# Unwinding of an RNA duplex by the Orthoflavivirus NS3 helicase requires translocation beyond the displaced strand and is stimulated by the NS5 RdRp

**DOI:** 10.64898/2026.02.06.704446

**Authors:** Jamie J. Arnold, Shubeena Chib, Craig E. Cameron

## Abstract

The NS3 helicases from the *Flaviviridae* family of viruses exhibit nucleotide-hydrolysis-dependent, nucleic-acid-unwinding activity. The RNA unwinding activity for NS3 helicases from the *Ortho*f*lavivirus* genus has not been fully explored and contrasts with NS3 helicase from Hepatitis C virus (HCV) of the *Hepacivirus* genus, which has thus far served as the prototypical model enzyme from this family of viruses. To begin to understand the functional differences between flavivirus NS3 helicases, we first developed an expression and purification system for full-length untagged NS3 protein from West Nile virus (WNV) and Zika virus (ZIKV). Both enzymes exhibit RNA-stimulated ATPase activity and are dependent on the nucleoside triphosphatase active site of the enzyme. Unlike HCV NS3, orthoflavivirus NS3s do not efficiently pre-assemble on a 3’-ssRNA-tailed dsRNA substrate in the absence of ATP-Mg which is a prerequisite for formation of a productive HCV NS3-RNA complex that can exhibit a rapid burst of RNA unwinding. Instead, to observe RNA unwinding by WNV and ZIKV NS3s, low Mg-ATP concentrations are required at a time coincident when NS3 encounters the RNA substrate. In addition, we find that orthoflavivirus NS3s require translocation beyond the displaced strand to completely unwind a dsRNA substrate. Last, we find that orthoflavivirus NS5 stimulates the ability of NS3 to unwind dsRNA. These results suggest that functional differences exist between the flavivirus NS3 helicases and illuminate that orthoflavivirus NS3s require a functional interaction with the NS5 protein for coordination of its activity, as it is believed these two proteins constitute the viral replicase.

## Introduction

Replication of positive-strand RNA viruses requires a coordinated series of enzymatic reactions catalyzed by viral enzymes assembled into membrane-associated replication complexes (1–5). For the orthoflaviviruses within the *Flaviviridae* family of viruses, viral RNA replication depends primarily on two enzymes: non-structural protein 5 (NS5), which contains an N-terminal methyltransferase and C-terminal RNA-dependent RNA polymerase (RdRp) domain, and non-structural protein 3 (NS3), which contains an N-terminal protease and C-terminal helicase domain (6–9). Together, these enzymes are thought to interact functionally to synthesize, modify, and unwind viral RNA during viral replication (10, 11). Understanding how these enzymes function individually and in concert is essential to define the molecular basis of orthoflavivirus RNA replication.

Helicases couple NTP hydrolysis to translocation and strand separation of nucleic acids (12–14). The NS3 helicase of *Flaviviridae* belongs to superfamily 2 (SF2) (15, 16) and has been studied most extensively in Hepatitis C virus (HCV) of the *Hepacivirus* genus, which has served as the prototype for this family (13, 17, 18). HCV NS3 exhibits robust, RNA-stimulated ATPase activity and catalyzes duplex RNA unwinding through formation of a pre-assembled enzyme–substrate complex in the absence of ATP–Mg²⁺ (19). Upon nucleotide addition, this complex undergoes a rapid burst of strand displacement (19). Whether the NS3 helicases from the *Orthoflavivirus* genus employ a similar mechanism has not been well established.

Despite high structural and sequence similarity (20), orthoflavivirus NS3 helicases may differ functionally from the HCV enzyme. Orthoflavivirus replication occurs exclusively in cytoplasmic replication organelles where NS3 acts in conjunction with NS5 (1–5, 10, 11). Structural studies have defined conserved catalytic motifs and overall fold (20), yet kinetic data describing RNA binding, ATP utilization, and strand displacement remain limited (20–39). Moreover, whether NS5 modulates NS3 activity directly has only recently been evaluated (10, 11, 38–45).

Here, we sought to define the biochemical properties of the orthoflavivirus NS3 helicases and to test whether NS5 influences their function. We developed bacterial expression and purification systems for full-length, untagged NS3 enzymes from West Nile virus (WNV) and Zika virus (ZIKV) and characterized their ATPase and helicase activities under a range of conditions. We show that RNA stimulates ATP hydrolysis by both enzymes, but productive unwinding requires low ATP–Mg²⁺ concentrations at the time of RNA engagement. Unlike HCV NS3, the orthoflavivirus enzymes fail to form stable pre-initiation complexes in the absence of nucleotide. We also demonstrate that NS3 prefers duplex substrates with a 5′-single-stranded tail and that the cognate NS5 polymerase enhances NS3-catalyzed strand separation. Together, these results define key mechanistic differences between the helicases of *Orthoflavivirus* genera and reveal that the NS3–NS5 interaction contributes directly to RNA unwinding.

## Results

### RNA stimulates the ATPase activity of WNV NS3

To determine whether WNV NS3 exhibits RNA-dependent ATPase activity, we first purified both wild-type (WT) WNV NS3 and a derivative in which the catalytic aspartic acid residue (Asp-285) was mutated to alanine (D285A NS3) (**Fig. 1A**). The proteins were expressed and purified as HIS-SUMO-NS3 fusion proteins and subsequently processed with the SUMO protease, Ulp1 to produce a full-length untagged NS3 protein. WT NS3 hydrolyzed ATP and was significantly stimulated in the presence of double-stranded RNA (ds9C20) (**Fig. 1B-D**). However, D285A NS3 did not show any demonstrable ATPase activity (**Fig. 1C**). The activity of WT NS3 increased from 180 ± 10 to 440 ± 10 µM/min/µM upon RNA addition (**Fig. 1D**). This RNA-mediated stimulation mirrors observations with the HCV NS3 ATPase, suggesting a conserved allosteric activation mechanism.

**Figure 1.**
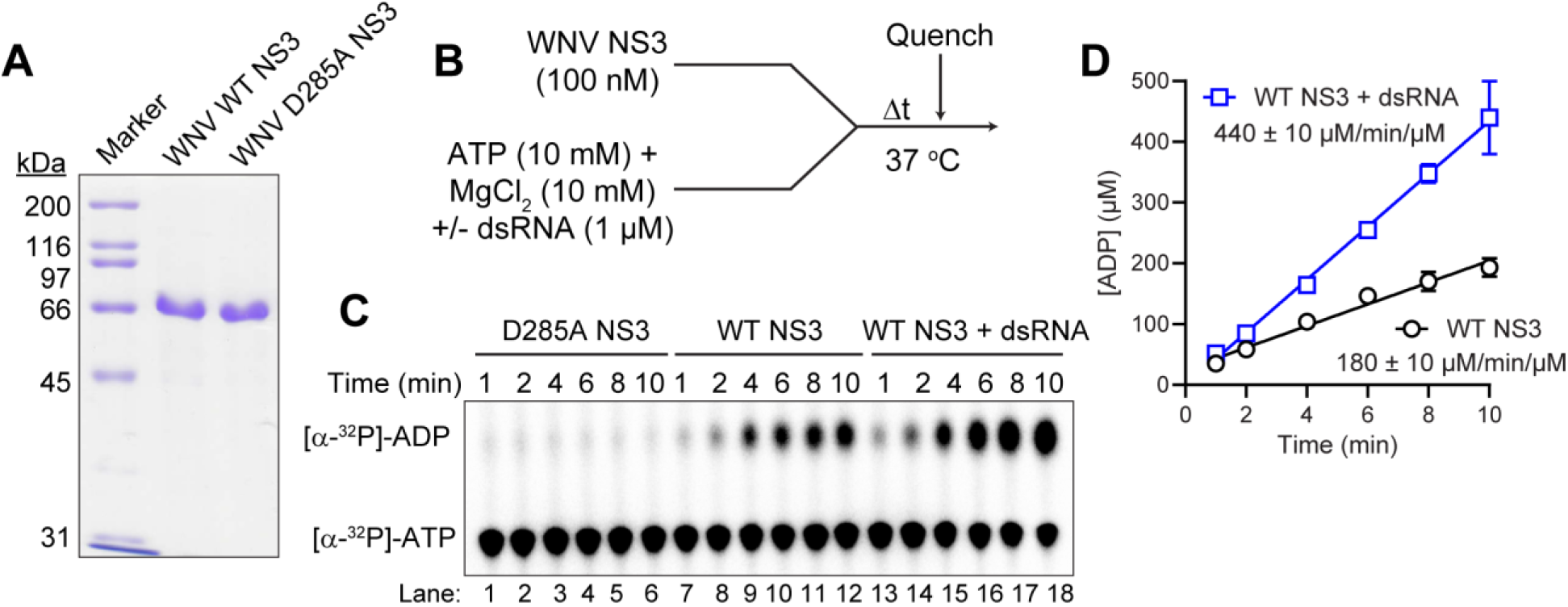
WNV NS3 shows RNA-stimulated ATPase activity. **(A)** 10% SDS-PAGE gel showing pure WNV WT NS3 and D285A NS3. 2 µg of protein was loaded on the gel. **(B)** A scheme showing the experimental set-up for NS3-catalyzed ATPase activity. **(C)** Representative TLC of NS3-catalyzed ATPase activity in the absence and presence of dsRNA. **(D)** Data were fit to a line yielding a specific activity of 180 ± 10 µM/min/µM and 440 ± 10 µM/min/µM in the absence and presence of RNA, respectively. Plotted are the means and SD from three replicates.

### WNV NS3 fails to unwind dsRNA under conditions optimized for HCV NS3

Despite robust ATPase activity, WNV NS3 was unable to unwind a model duplex RNA substrate (ds9C20) under conditions previously optimized for HCV NS3 (1 µM enzyme, 10 mM ATP/MgCl₂, 37 °C) (**Fig. 2**) (19). In contrast, HCV NS3 catalyzed efficient strand separation under identical conditions. Reducing the reaction temperature did not result in any measurable unwinding by WNV NS3 (**Fig. 2C**). These findings suggest that WNV NS3 either requires distinct reaction conditions or a different substrate architecture to promote strand displacement.

**Figure 2.**
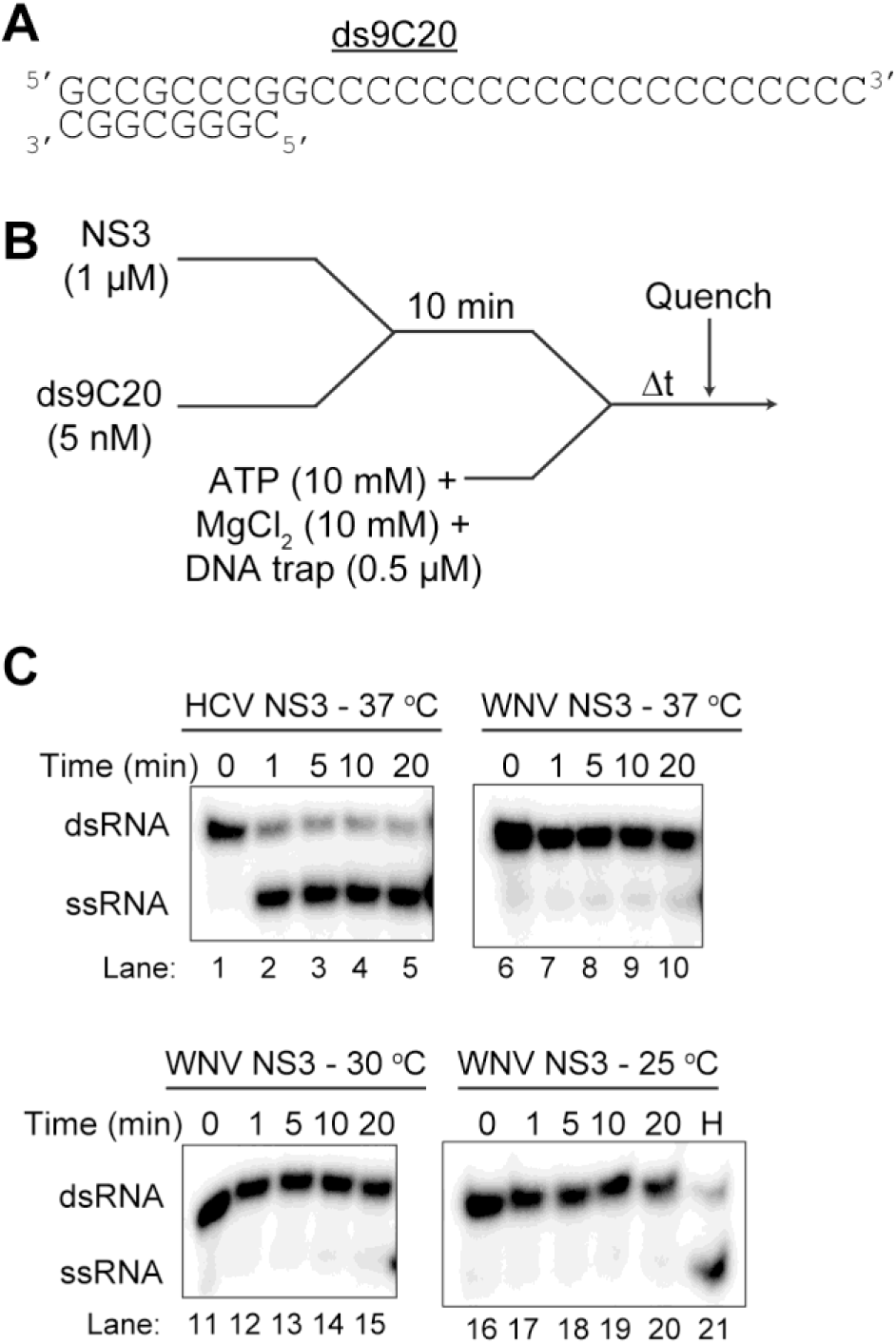
WNV NS3 does not unwind dsRNA substrate under conditions optimized for HCV NS3. **(A)** HCV NS3 model unwinding substrate (dsRNA) that consists of a 9 bp duplex region and 20 nt ss-overhang on the 3’ end. **(B)** A scheme showing the experimental set-up for NS3-catalyzed unwinding of ds9C20. WNV or HCV NS3 (1 µM) was pre-incubated with ds9C20 RNA substrate (5 nM) at either 25 or 30 or 37⁰C for 10 minutes. Unwinding was initiated by addition of ATP (10 mM), MgCl_2_ (10 mM) and DNA trap (0.5 µM). The reactions were quenched at increasing times. **(C)** Products of the reactions were resolved from substrates by 20% native PAGE and visualized by phosphorimaging. Representative phosphorimages are shown. H is heat control.

### WNV NS3-mediated RNA unwinding requires lower ATP and MgCl₂ concentrations

To test whether elevated ATP and MgCl_2_ concentrations impair unwinding, we performed helicase assays at reduced ATP (2 mM) and MgCl₂ (0.5 mM) concentrations (**Fig. 3**). These conditions are like previous studies on DENV NS3 (32). Under these modified conditions, WNV NS3 unwound ds9C20 RNA at 30 °C (**Fig. 3B**). WNV NS3 unwinding activity was lower at 37 °C (**Fig. 3B**). In contrast, HCV NS3 completely unwound the duplex at 37 °C but at 30 °C produced less ssRNA product (**Fig. 3B**). We performed a time course with WNV NS3 and fit the data to a single exponential yielding a rate constant of 0.22 ± 0.04 min⁻¹ and an amplitude of 32% (**Fig. 3C,D**). These results highlight the importance of optimizing cofactor concentrations to support unwinding activity by orthoflaviviral helicases.

**Figure 3.**
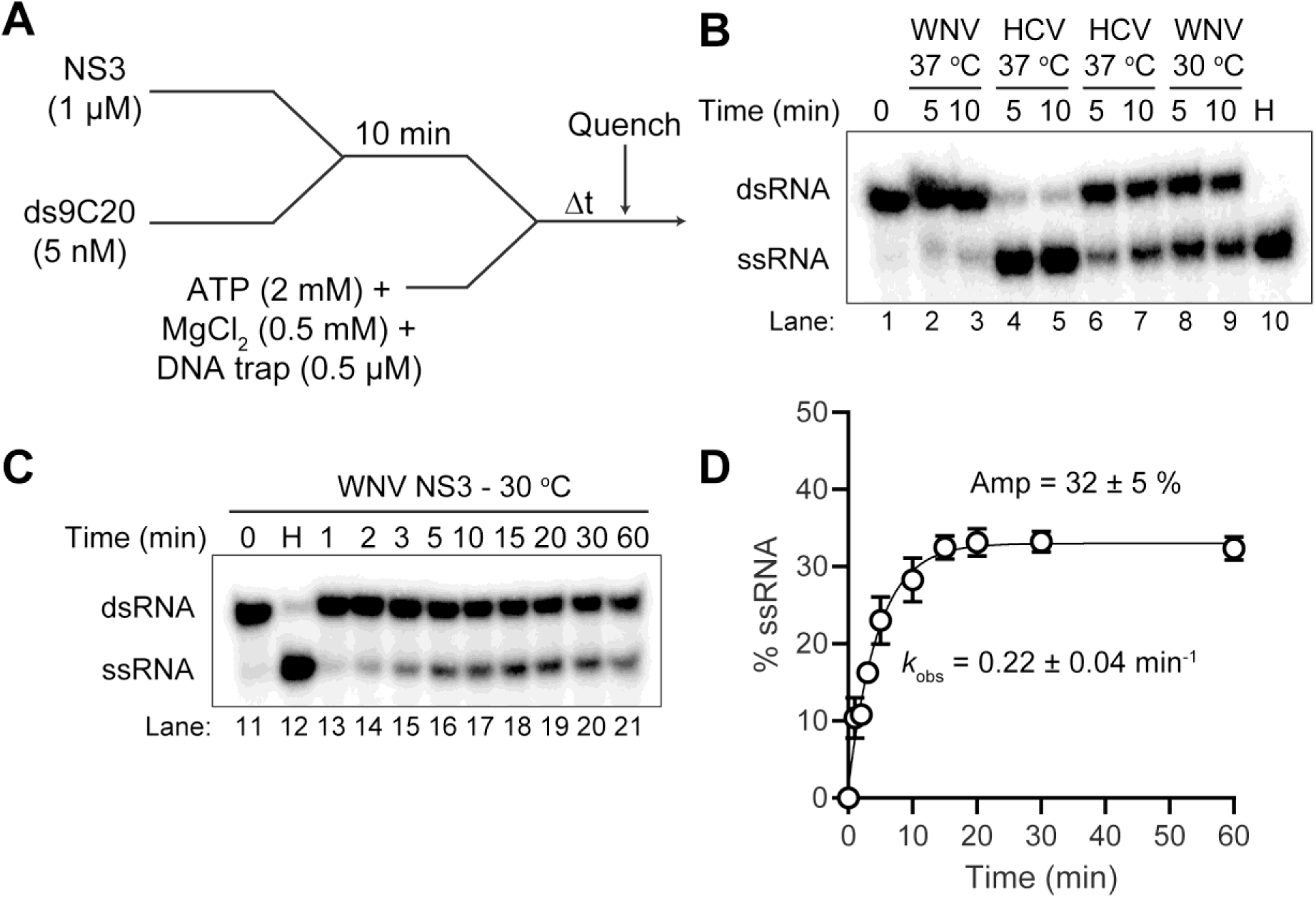
Unwinding of ds9C20 occurs by WNV NS3 with lower concentrations of ATP and MgCl_2_. **(A)** A schematic representation of the experimental set-up for WNV NS3-catalyzed unwinding of ds9C20. WNV or HCV NS3 (1 µM) was pre-incubated with ds9C20 RNA substrate (5 nM) at either 30 or 37⁰C for 10 minutes. Unwinding was initiated by addition of ATP (2 mM), MgCl_2_ (0.5 mM) and DNA trap (0.5 µM). The reactions were quenched at increasing times by addition of EDTA (200 mM) and SDS (0.33%). Products of the reactions were resolved from substrates by 20% native PAGE and visualized by phosphorimaging. **(B)** Representative gel image showing the unwinding of dsRNA by HCV or WNV NS3 at 30 or 37⁰C. **(C)** Gel image indicates a time course reaction for WNV NS3 at 30⁰C. H is heat control. **(D)** Phosphorimage was quantitated and %ssRNA produced was plotted as a function of reaction time. The data were fit to a single exponential, yielding a rate constant of 0.22 ± 0.04 min^-1^. Plotted are the means and SD from three replicates.

### RNA binding by WNV NS3 is sensitive to ATP and MgCl₂

We next assessed how ATP and MgCl₂ affect RNA binding by WNV NS3 (**Fig. 4** and **Table 1**). Fluorescence polarization assays revealed that the apparent dissociation constants (*K*d,app) for both single-stranded and double-stranded RNA substrates increased markedly in the presence of ATP/MgCl₂, indicating weakened binding affinity (**Table 1**). These data suggest that the transition from substrate binding to catalysis is coupled to ATP-induced conformational changes, as previously demonstrated for HCV NS3.

**Figure 4.**
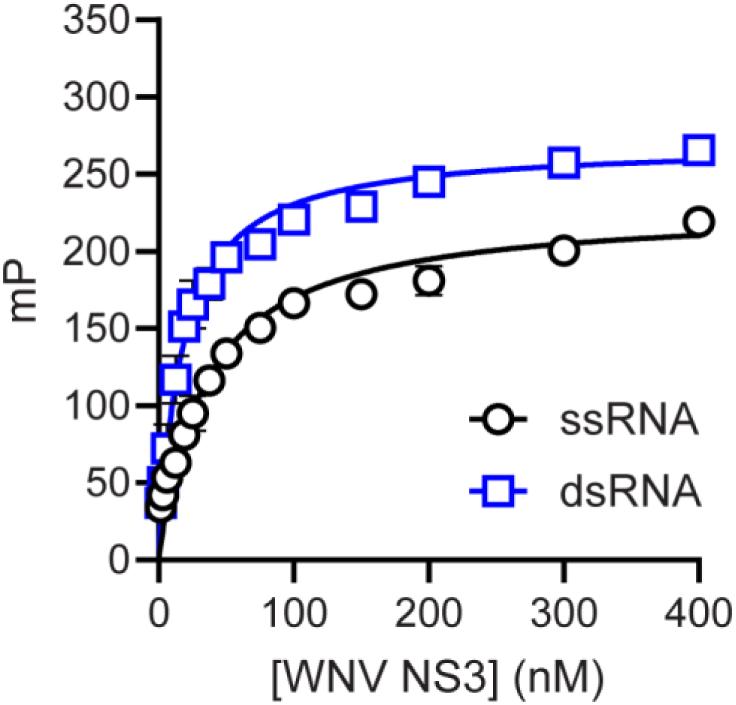
Affinity of WNV NS3 for ssRNA and dsRNA. Fluorescein-labeled ssRNA (ss9-C20) or dsRNA (ds9C20) were incubated with increasing concentrations of WNV NS3 for 1 minute. The change in fluorescence millipolarization (mP) was subsequently monitored. mP values were plotted as a function of NS3 concentrations and the data were fit to a hyperbola. A representative binding isotherm is shown for the interaction of NS3 with ssRNA and dsRNA in the absence of ATP-MgCl_2_. Data were fit to a hyperbola yielding an apparent dissociation constant (*K*d,app) of 62 ± 8 nM and 22 ± 3 nM for ssRNA and dsRNA, respectively. Plotted are the means and SD from three replicates.

**Table 1.**
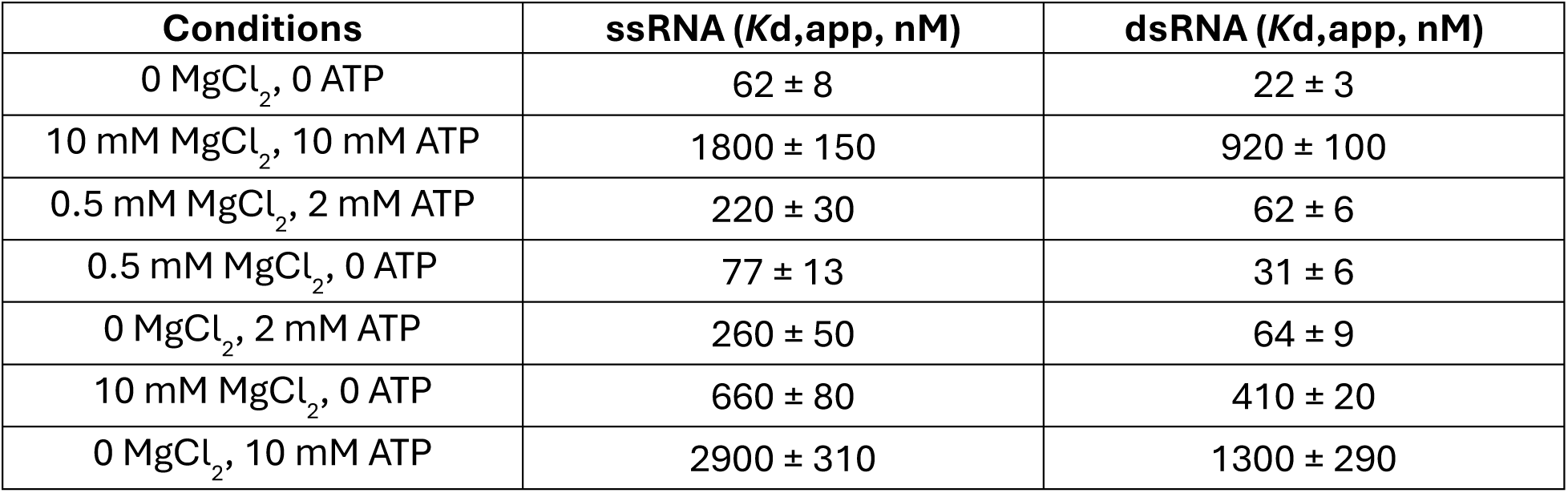
Affinity of WNV NS3 for ssRNA and dsRNA decreases with high concentrations of ATP and MgCl_2_. Apparent dissociation constants for ssRNA (ss9C20) or dsRNA (ds9C20) for WNV NS3 under various conditions of MgCl_2_ and ATP concentrations. Fluorescein-labeled ssRNA or dsRNA was mixed with increasing concentrations of WNV NS3 under the various conditions reported. Millipolarization (mP) was plotted and the data were fit to a hyperbola as shown in Figure 4, yielding the apparent dissociation constants (*K*d,app). Values are rounded to two significant figures.

### Optimal helicase activity requires NS3 concentration above the *K*_d_ for RNA binding

We titrated the NS3 concentration in RNA unwinding assays to determine the minimal enzyme requirement for maximal helicase activity (**Fig. 5**). Strand separation plateaued at ∼700 nM NS3, consistent with a concentration approximately 5-10-fold above the RNA-binding *K*d,app using 0. 5 mM MgCl_2_ and 2 mM ATP (**Fig. 5B,C**). Higher concentrations became inhibitory. These data indicate that a high enzyme-to-substrate ratio is critical for promoting unwinding by WNV NS3.

**Figure 5.**
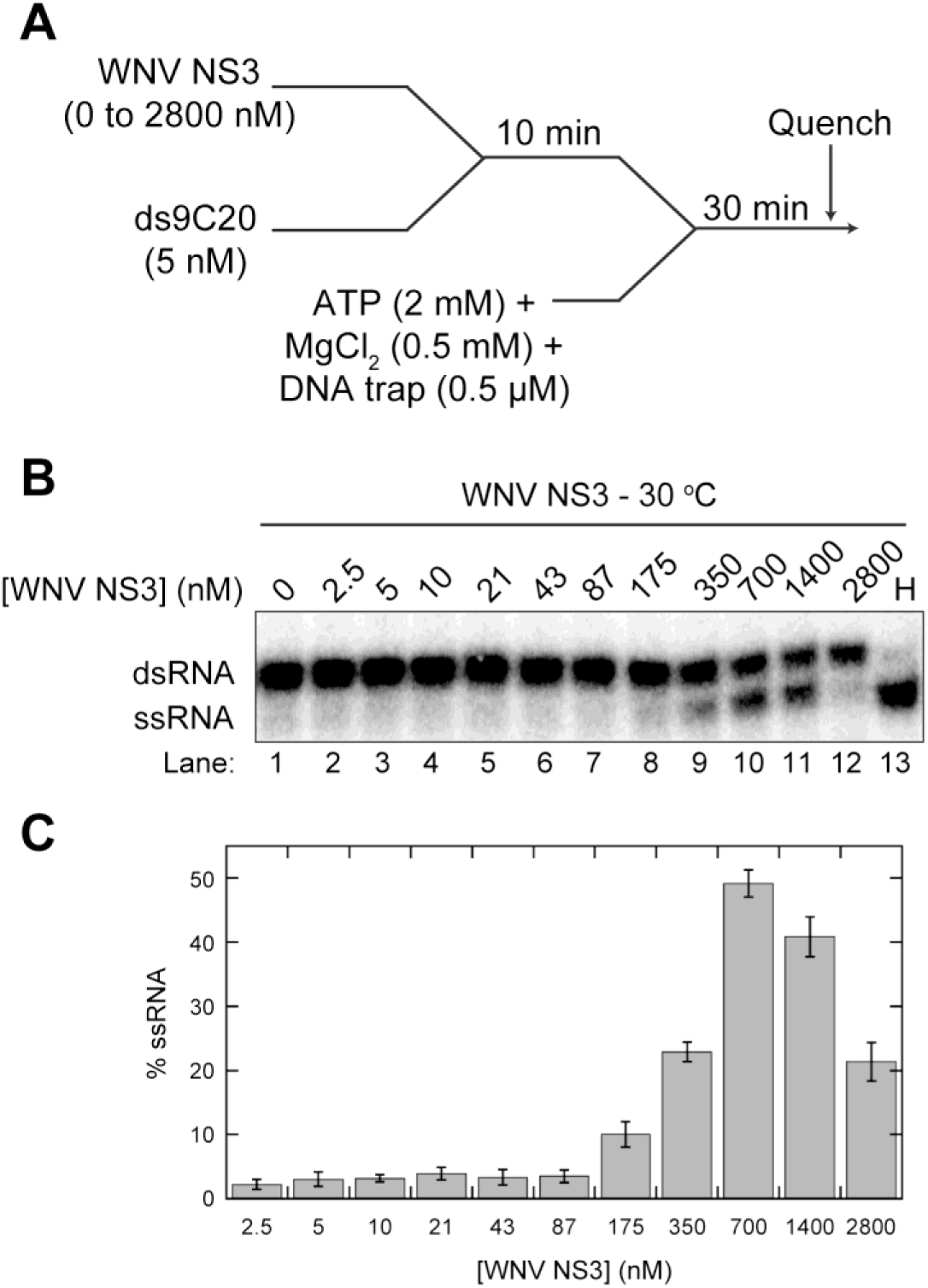
NS3 concentration required for optimal unwinding is consistent with being 10x above the *K*d,app for unwinding substrate. **(A)** A schematic representation of the experimental set-up for WNV NS3-catalyzed unwinding of ds9C20. Varying concentrations of WNV NS3 was pre-incubated with ds9C20 RNA substrate (5 nM) at 30 ⁰C for 10 minutes. Unwinding was initiated by addition of ATP (2 mM), MgCl_2_ (0.5 mM) and DNA trap (0.5 µM). The reactions were quenched at 30 minutes. Products of the reactions were resolved from substrates by 20% native PAGE and visualized by phosphorimaging. **(B)** Representative gel image indicating product formation in the presence of varying enzyme concentrations. H is heat control. **(C)** The phosphorimage was quantified by ImageQuant software and %ssRNA formed was plotted as a function of NS3 concentration. Plotted are the means and SD from three replicates.

### WNV NS3 does not efficiently form a stable pre-assembled complex with dsRNA substrate in the absence of ATP/MgCl_2_

To probe whether NS3 pre-binds RNA before catalysis and unwinding, we varied the pre-incubation time of NS3 with dsRNA substrate prior to ATP/MgCl₂ addition (**Fig. 6**). No significant increase in unwinding efficiency was observed with longer pre-incubation times, indicating that NS3-dsRNA complex formation is not in the most competent state for rapid unwinding in the absence of ATP (**Fig. 6B,C**). This stands in contrast to HCV NS3, which preassembles onto the dsRNA substrate in the absence of ATP/MgCl_2_ and exhibits rapid unwinding activity once the reaction is initiated with ATP/MgCl_2_.

**Figure 6.**
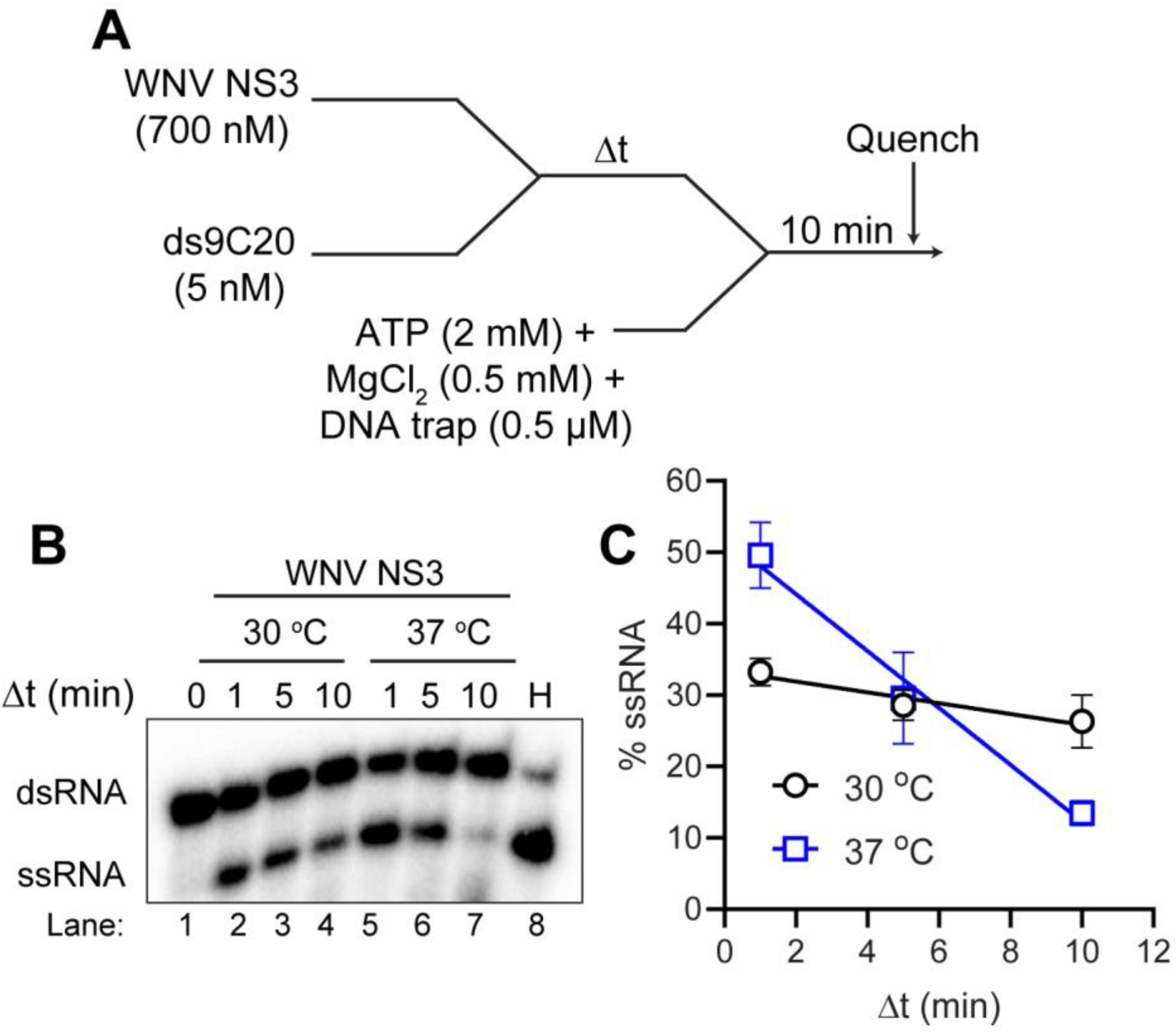
The WNV NS3-RNA complex does not pre-assemble prior to the addition of ATP-MgCl_2_. **(A)** A scheme showing the experimental set-up for NS3-catalyzed unwinding of ds9C20. WNV NS3 (700 nM) was pre-incubated with ds9C20 RNA substrate (5 nM) at either 30 or 37⁰C for varying amounts of time (1, 5, and 10 minutes). Unwinding was initiated by addition of ATP (2 mM), MgCl_2_ (0.5 mM) and DNA trap (0.5 µM). Each reaction was quenched at 10 minutes. Products of the reactions were resolved from substrates by 20% native PAGE and visualized by phosphorimaging. **(B)** Representative gel image indicating product formation in the presence of varying pre-incubation time at 30 and 37 ⁰C. H is heat control. **(C)** The phosphorimage was quantified by ImageQuant software and %ssRNA produced was plotted as a function of pre-incubation time using Kaleidagraph software. Plotted are the means and SD from three replicates.

### Strand separation by NS3 is rapid and requires ATP hydrolysis

To directly assess the ATP dependence of helicase activity, we compared WNV WT NS3 to D285A NS3 in RNA unwinding assays; HCV NS3 was used as a control (**Fig. 7**). Only the WNV WT NS3 enzyme promoted strand separation, no product was observed with D285A NS3, with a rate constant of 0.27 ± 0.10 min⁻¹ (**Fig. 7D**). In the absence of ATP, WNV WT NS3 did not promote unwinding. These data confirm that unwinding activity is strictly dependent on ATP hydrolysis, as expected for a canonical SF2 helicase mechanism.

**Figure 7.**
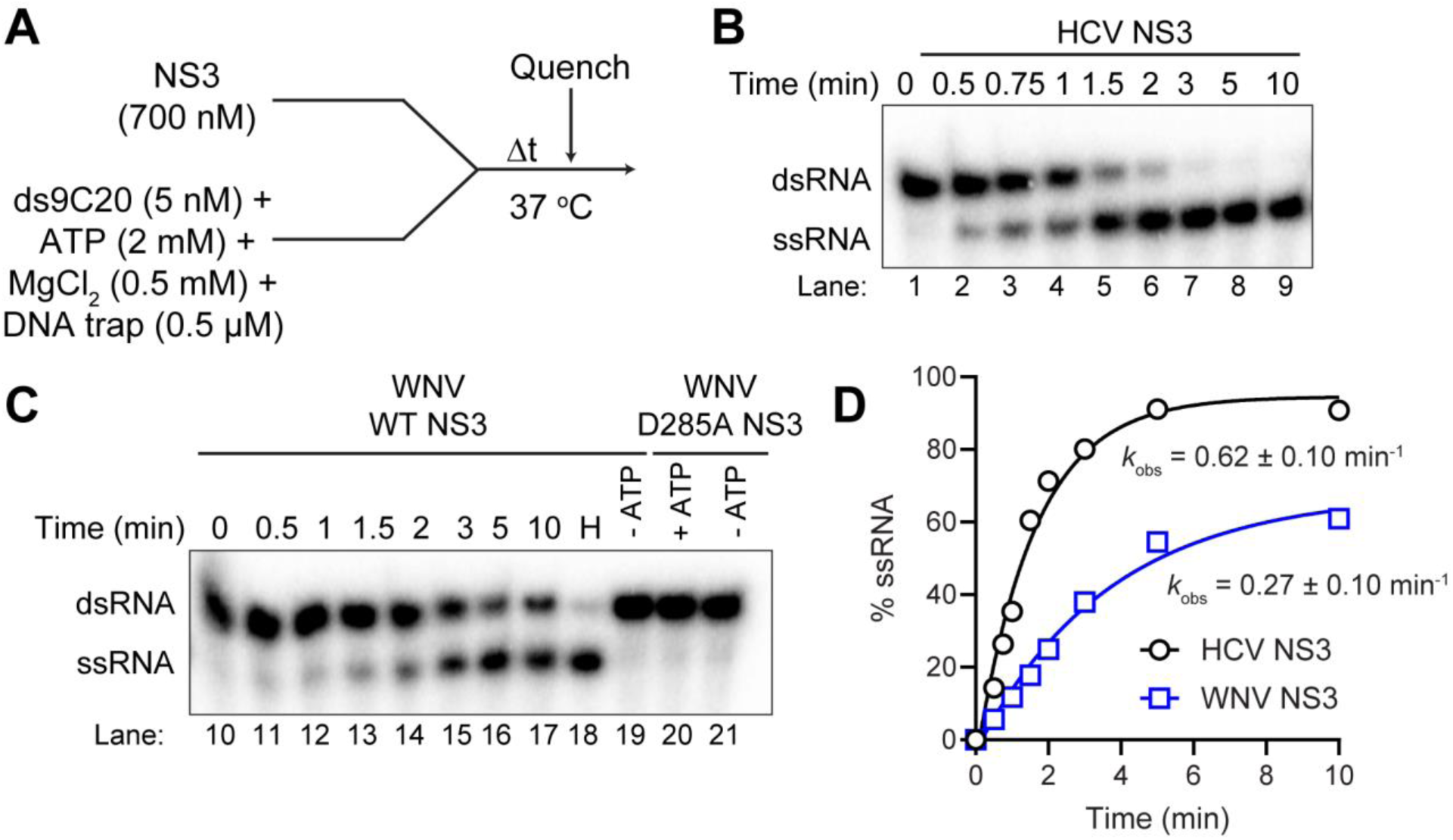
Initiating reaction with WNV NS3 leads to rapid unwinding of dsRNA which is dependent on ATP hydrolysis. **(A)** A scheme representing the experimental set-up for NS3-catalyzed unwinding of ds9C20. A master mix containing dsRNA substrate ds9C20 (5 nM), ATP (2 mM), MgCl_2_ (0.5 mM) and DNA trap (0.5 µM) was prepared. Unwinding reaction was initiated at 37⁰C by addition of either HCV NS3 (700 nM), WNV WT NS3 (700 nM), or WNV ATPase-dead D285A NS3 (700 nM). Reactions were quenched at increasing times. Products of the reactions were resolved from substrates by 20% native PAGE and visualized by phosphorimaging. **(B)** Representative gel image showing the unwinding of dsRNA by HCV NS3. **(C)** Representative phosphorimage is shown for WNV WT NS3 and D285A NS3. Lanes 19 and 20 are performed in the absence of ATP after 10 min reaction time. H is heat control. **(D)** Phosphorimages were quantitated and %ssRNA produced was plotted as a function of reaction time. The data were fit to a single exponential, yielding a rate constant of 0.27 ± 0.10 min^-1^ and 0.62 ± 0.10 min^-1^ for WNV NS3 and HCV NS3, respectively.

### WNV NS3 exhibits complete dsRNA unwinding activity with a substrate that has a 5’-ssRNA tail beyond the duplex region

We next tested whether altering RNA substrate architecture could enhance NS3 activity. WNV NS3 translocates in a 3’-to-5’ direction and therefore requires a 3’-ssRNA loading strand. We reasoned that extending ssRNA beyond the length of the duplex might allow the enzyme to remain stably bound to the substrate and completely separate the duplex (**Fig. 8A,B**). A modified RNA duplex (C10ds9C20) featuring a 5′-single-stranded tail on the loading strand supported more efficient unwinding than the original ds9C20 substrate (**Fig. 8C**). The observed rate constant for strand separation increased to 0.59 ± 0.10 min⁻¹ versus 0.27 ± 0.10 min⁻¹ for the untailed duplex and reached near 100% unwinding (**Fig. 8C,D**). These results suggest that WNV NS3 has an intrinsic preference for substrates with defined polarity and loading strand architecture; needing to remain stably bound to the RNA substrate to facilitate complete unwinding.

**Figure 8.**
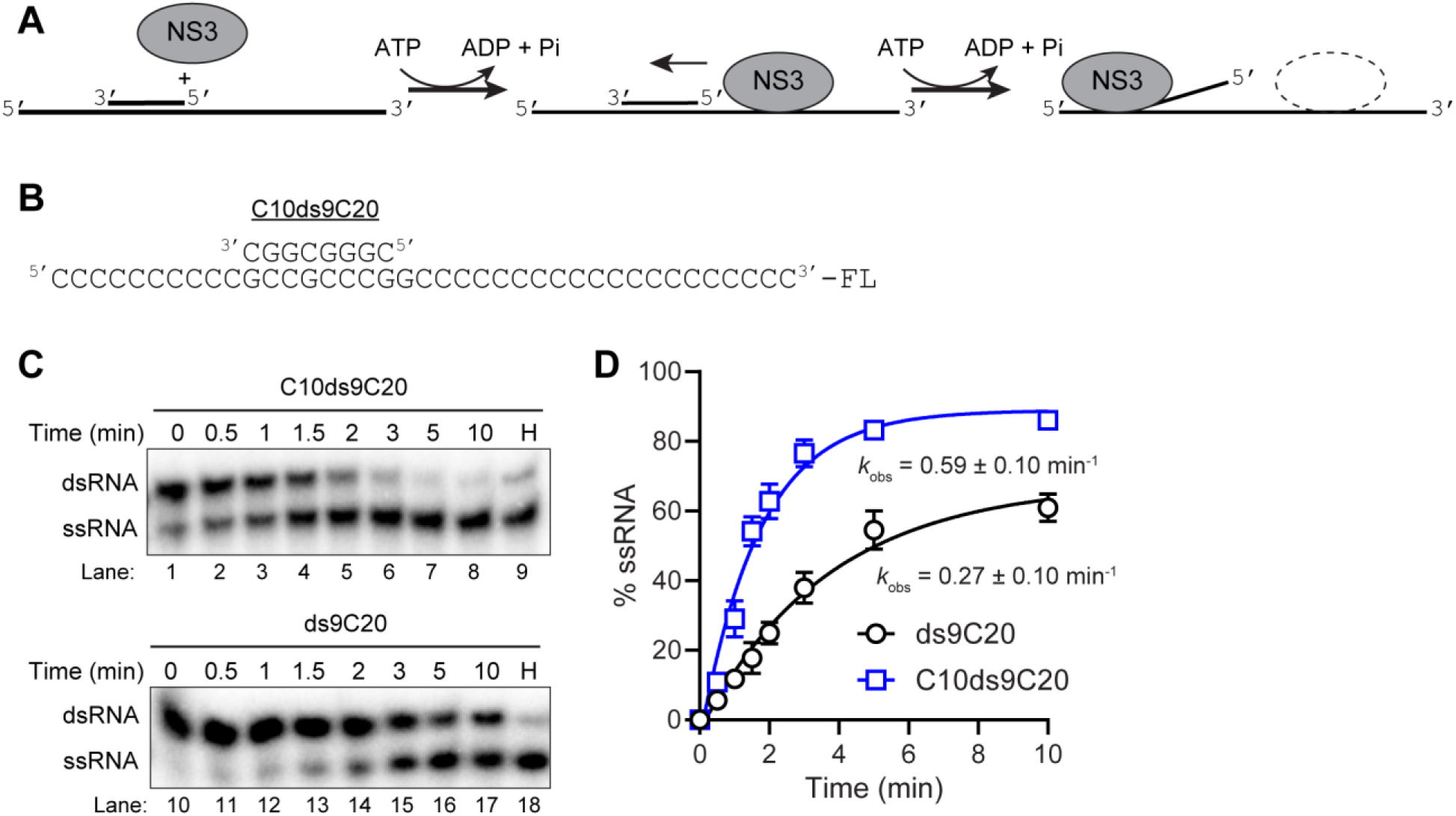
WNV NS3 shows preference for unwinding a substrate that has a 5’-ssRNA tail on the loading strand. **(A,B)** A scheme showing NS3-catalyzed unwinding of C10ds9C20 (panel B), an RNA substrate that has an additional single-stranded overhang on the 5’-end of the loading strand. A master mix containing C10ds9C20 (5 nM), ATP (2 mM), MgCl_2_ (0.5 mM) and DNA trap (0.5 µM) was prepared. Unwinding reaction was initiated at 37⁰C by addition of WNV WT NS3 (700 nM). Reactions were quenched at increasing times. As a control assay, the model substrate ds9C20 was also unwound alongside under the same conditions. Products of the reactions were resolved from substrates by 20% native PAGE and visualized by phosphorimaging. **(C)** Representative phosphorimages show the time course unwinding assay for C10ds9C20 and ds9C20 dsRNA substrates. H is heat control. **(D)** The images were quantified and %ssRNA produced was plotted as a function of reaction time. The data were fit to a single exponential, yielding a rate constant of 0.59 ± 0.10 min^-1^ and 0.27 ± 0.10 min^-1^ for C10ds9C20 and ds9C20, respectively. Plotted are the means and SD from three replicates.

### WNV and ZIKV NS3 display comparable ATPase and helicase activities under identical conditions

To assess whether these observations generalize across orthoflaviviruses, we evaluated ATPase and helicase activity of ZIKV NS3 under optimized conditions (**Fig. 9**). ZIKV NS3 hydrolyzed ATP with basal and RNA-stimulated specific activities of 180 ± 10 and 460 ± 20 µM/min/µM, respectively. This translated to robust helicase activity with a rate constant of 1.4 ± 0.06 min⁻¹ for the C10ds9C20 substrate (**Fig. 9F,G**), demonstrating that the biochemical behavior of ZIKV NS3 closely parallels that of its WNV ortholog.

**Figure 9.**
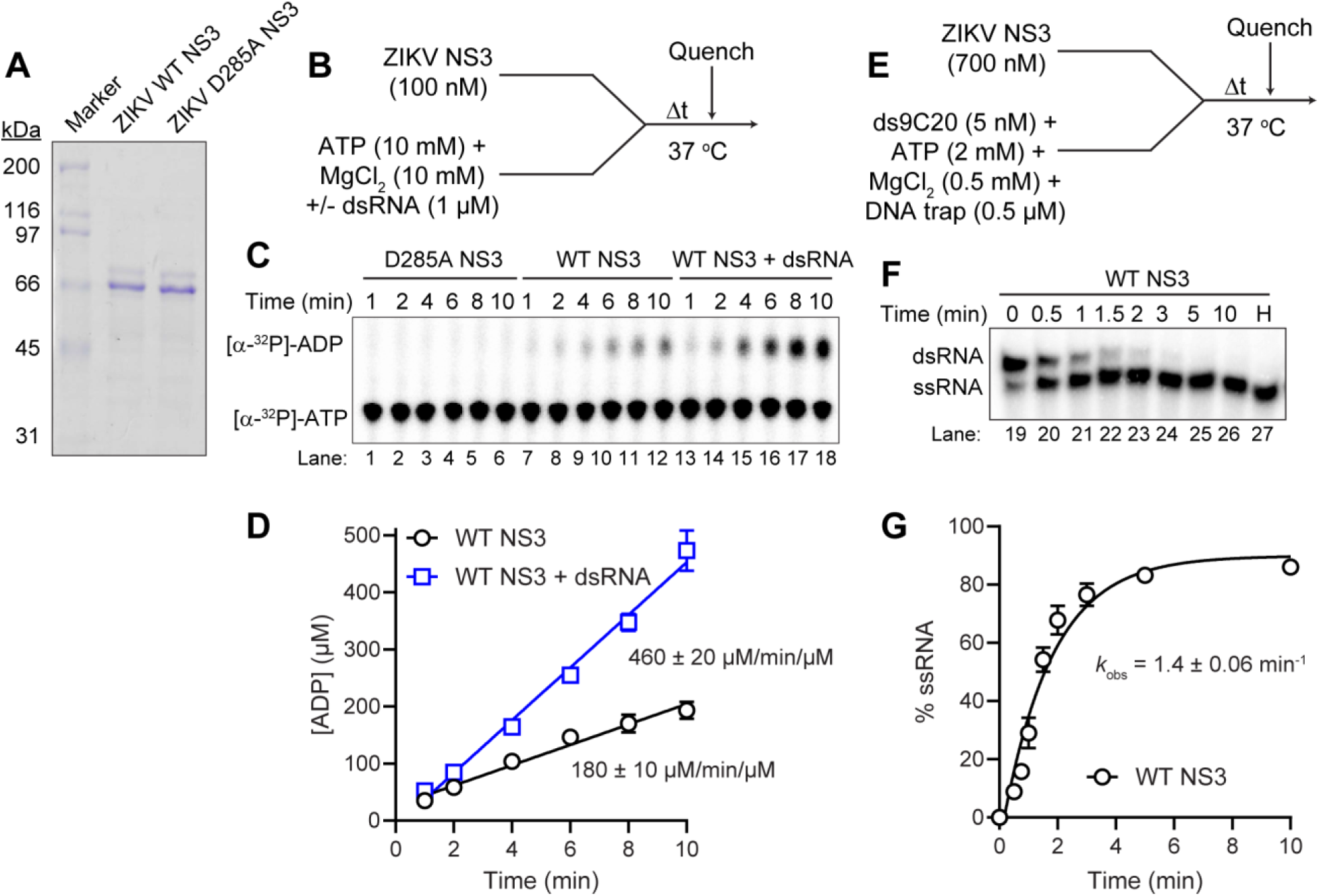
ATPase and helicase activities of ZIKV NS3. **(A)** 10% SDS-PAGE gel showing pure ZIKV WT NS3 and its ATPase-deficient mutant, D285A NS3. **(B)** Schematic of the experiment showing RNA-stimulated ATP hydrolysis activity of ZIKV NS3. NS3 (100 nM) was incubated with a mixture of [α^32^P]-ATP (10 mM) and MgCl_2_ (10 mM) in the absence or presence of ds9C20 (1 µM) at 37 ⁰C for 10 minutes. The reaction was quenched at various times by addition of EDTA (50 mM). **(C)** Product (ADP) was resolved from substrate (ATP) by thin layer chromatography (TLC), visualized by phosphorimaging, and quantified by ImageQuant software. **(D)** The ATP hydrolysis data were fit to a line yielding a specific activity of 180 ± 10 µM/min/µM and 460 ± 20 µM/min/µM in the absence and presence of RNA, respectively. Plotted are the means and SD from three replicates. **(E)** A scheme representing the experimental set-up for ZIKV NS3-catalyzed unwinding of dsRNA. A master mix containing dsRNA substrate C10ds9C20 (5 nM), ATP (2 mM), MgCl_2_ (0.5 mM) and DNA trap (0.5 µM) was prepared. Unwinding reaction was initiated at 37⁰C by addition of ZIKV WT NS3 (700 nM). Reactions were quenched at increasing times. **(F)** Products of the unwinding reaction were resolved from substrates by 20% native PAGE and visualized by phosphorimaging. Representative gel image shows the time course assay. H is heat control. **(G)** The image was quantified and %ssRNA produced was plotted as a function of time. The data were fit to a single exponential, yielding a rate constant of 1.4 ± 0.06 min^-1^ for C10ds9C20. Plotted are the means and SD from three replicates.

### Orthoflavivirus NS5 increases the efficiency of orthoflavivirus NS3 helicase activity

Because orthoflavivirus genome replication occurs through coordinated action of the NS3 helicase and the NS5 polymerase, we asked whether NS5 might modulate the RNA unwinding activity of NS3. To test whether the orthoflavivirus NS5 polymerase influences the helicase activity of NS3, we performed unwinding assays in the presence or absence of purified recombinant NS5 protein (**Fig. 10**). When equimolar concentrations of ZIKV NS5 and NS3 were combined, a marked increase in RNA unwinding was observed compared to NS3 alone (**Fig. 10C**). To determine whether this enhancement was specific to cognate orthoflavivirus partners, we compared NS5 proteins from ZIKV, WNV, and DENV for their ability to stimulate ZIKV and WNV NS3 (**Fig. 11**). Each NS5 stimulated the activity of its cognate NS3. Heterologous combinations also showed stimulation. Addition of poliovirus (PV) 3Dpol, a non-orthoflaviviral polymerase control, did not affect unwinding by either NS3 (**Fig. 11**), confirming specificity of the NS3-NS5 functional interaction. These findings suggest that orthoflavivirus NS5 directly interacts with the NS3 helicase to promote efficient unwinding of duplex RNA substrates. Finally, to verify that NS5 does not possess RNA unwinding activity and that stimulation requires a catalytically active NS3, we repeated the experiments using the ZIKV, WNV, and DENV NS5 alone and the ATPase-defective ZIKV D285A NS3 derivative in the presence of NS5 (**Fig. 12**). We also evaluated HCV NS5B to stimulate ZIKV NS3 unwinding. Neither ZIKV, WNV, or DENV NS5 alone nor ZIKV D285A NS3 with ZIKV NS5 promoted RNA strand separation (**Fig. 12**), confirming that NS5 does not independently unwind dsRNA and that its effect depends on an active NS3 helicase. HCV NS5B also had no impact on ZIKV NS3 (**Fig.12**). Collectively, these results support a model in which NS5 binding enhances the stability or conformational state of NS3 on RNA, thereby facilitating productive dsRNA unwinding activity.

**Figure 10.**
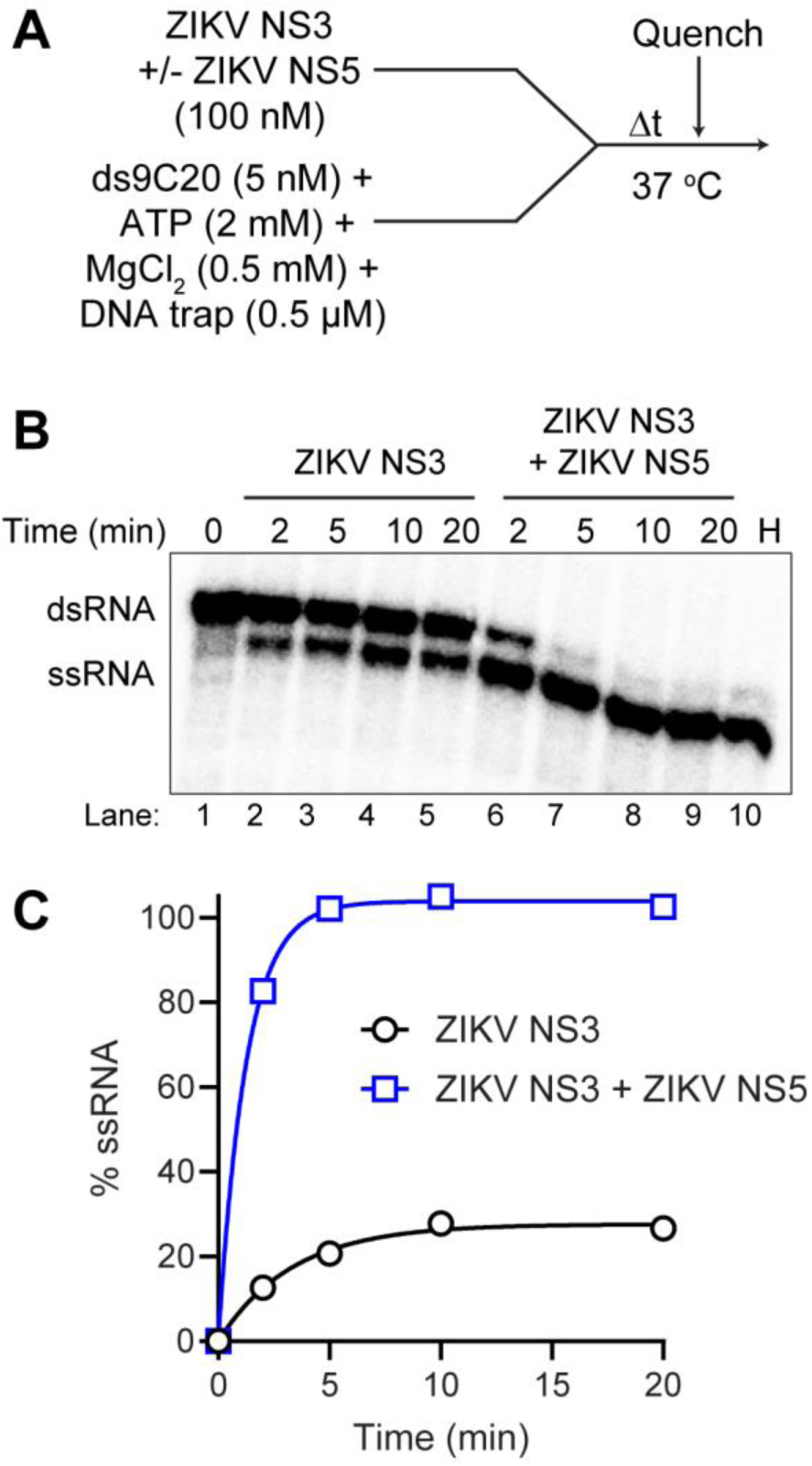
Orthoflavivirus NS5 increases the efficiency of orthoflavivirus NS3 helicase activity. **(A)** Schematic representation of the experimental setup for NS3-catalyzed unwinding of dsRNA in the absence or presence of NS5. Reactions contained 5 nM ds9C20 substrate, 2 mM ATP, 0.5 mM MgCl₂, 0.5 µM RNA trap and 100 nM ZIKV NS3 with or without 100 nM ZIKV NS5 final. ZIKV NS3 was pre-incubated alone or with ZIKV NS5 on ice prior to initiating the reaction. **(B)** Representative gel image showing the unwinding of dsRNA by ZIKV NS3 in the absence and presence of ZIKV NS5. H is heat control. **(C)** The image was quantified and %ssRNA produced was plotted as a function of time. The data were fit to a single exponential, yielding a rate constant of 0.30 ± 0.10 min^-1^ and 0.80 ± 0.10 min^-1^ and amplitude of 30% and 100% for ZIKV NS3 or ZIKV NS3+ZIKV NS5, respectively.

**Figure 11.**
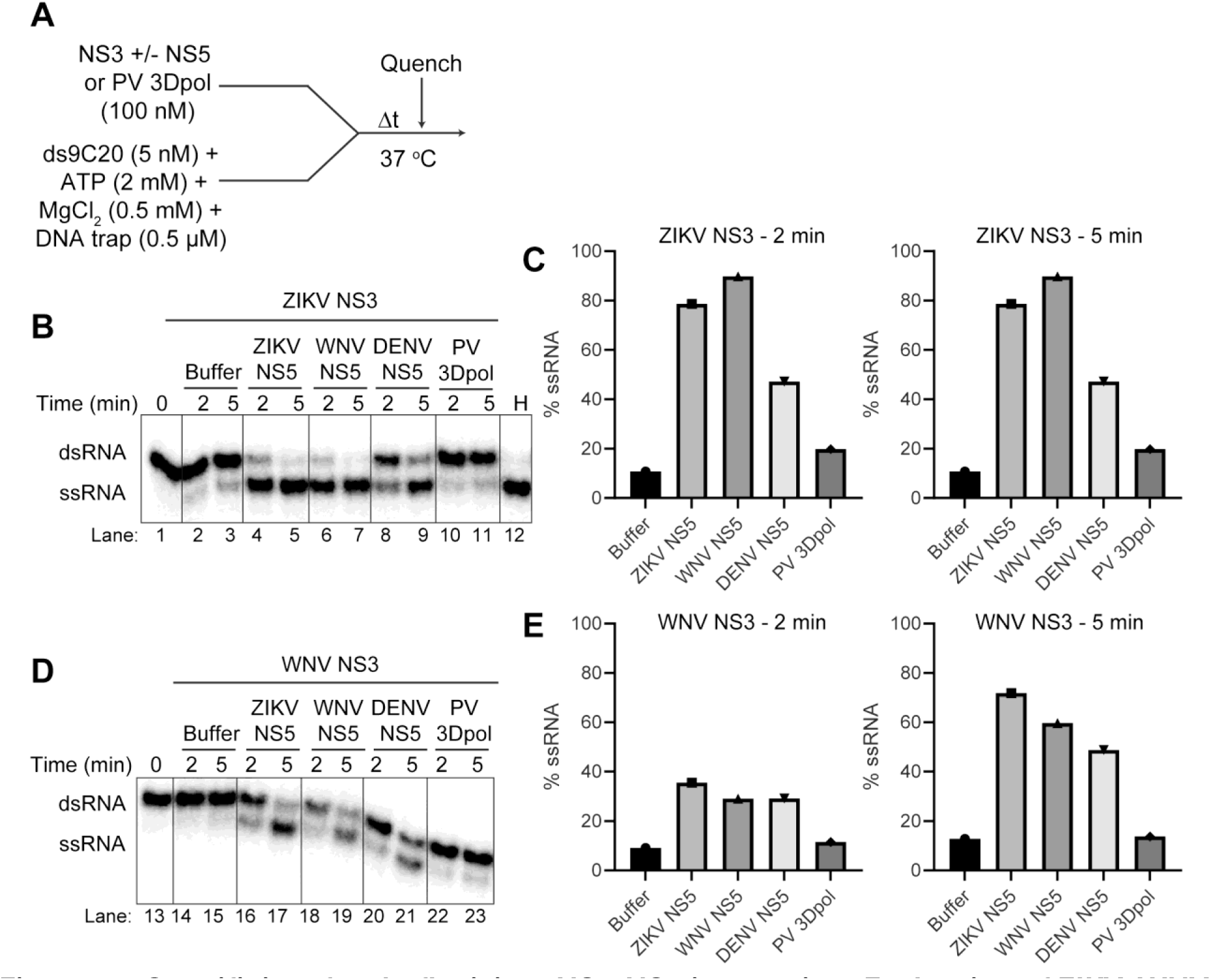
Specificity of orthoflavivirus NS3-NS5 interaction: Evaluation of ZIKV, WNV, DV NS5, and PV 3Dpol on ZIKV and WNV NS3 helicase activity. **(A)** Schematic representation of the experimental setup for NS3-catalyzed unwinding of dsRNA in the absence or presence of ZIKV, WNV, DENV NS5 or PV 3Dpol. Reactions contained 5 nM ds9C20 substrate, 2 mM ATP, 0.5 mM MgCl₂, 0.5 µM RNA trap and 100 nM NS3 with or without 100 nM NS5 (or PV 3Dpol) final. Either ZIKV NS3 or WNV NS3 was pre-incubated alone or with different flavivirus NS5’s or PV 3Dpol on ice prior to initiating the reaction. **(B-E)** Representative gel images (panels B and D) showing the unwinding of dsRNA by ZIKV NS3 or WNV NS3 in the absence and presence of NS5 from ZIKV, WNV, DENV, or PV 3Dpol. H is heat control. The phosphorimages were quantified by ImageQuant software and %ssRNA formed was plotted for both ZIKV or WNV NS3 in the absence and presence of NS5 from ZIKV, WNV, DENV, or PV 3Dpol.

**Figure 12.**
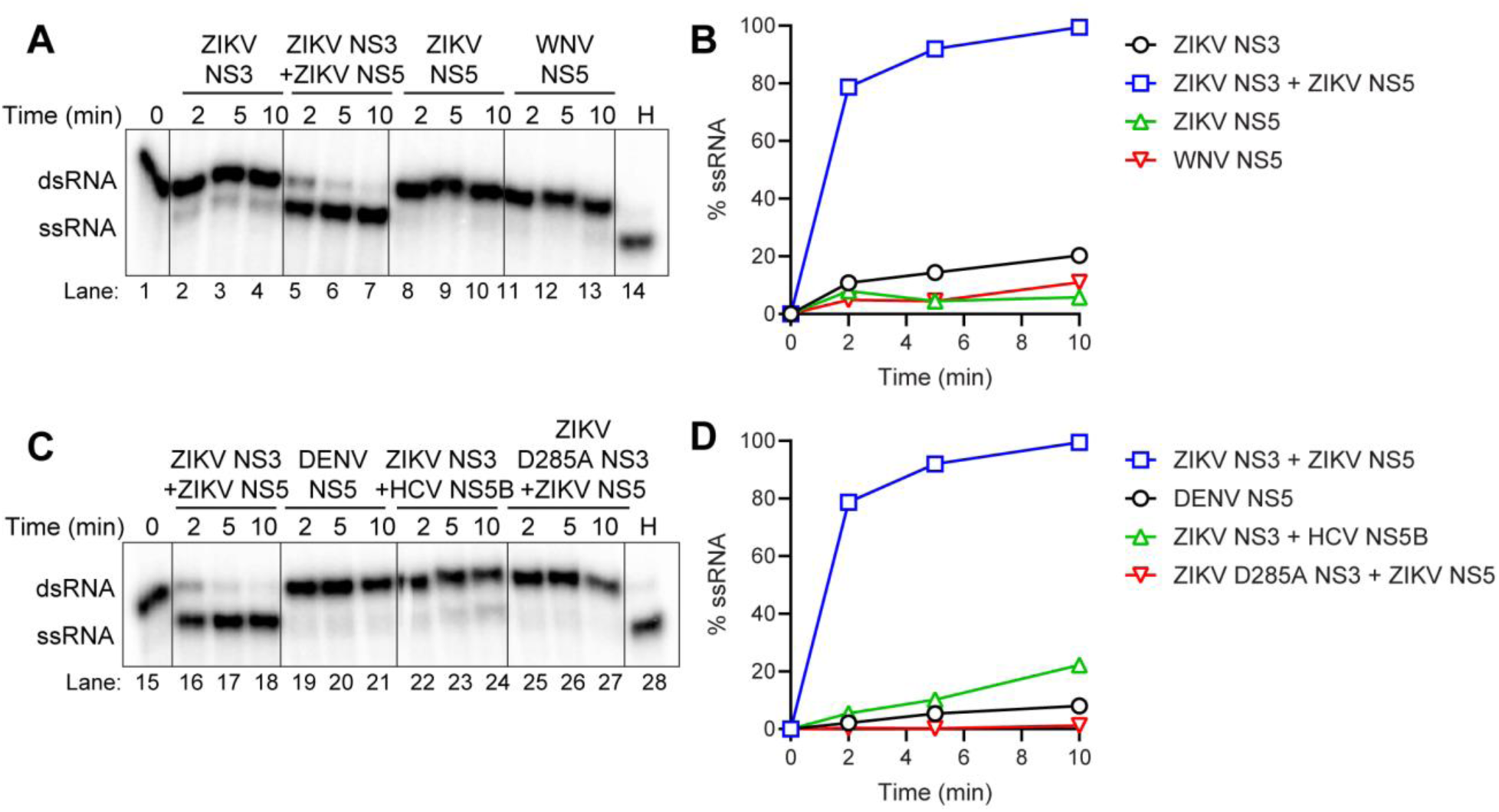
Neither Orthoflavivirus NS5 nor NS3 D285A in the presence of NS5 unwinds dsRNA. Reactions contained 5 nM ds9C20 substrate, 2 mM ATP, 0.5 mM MgCl₂, 0.5 µM RNA trap and either 100 nM WT NS3, D285A NS3, or NS5 final. **(A-D)** Representative gel images (panel A and C) showing the unwinding of dsRNA for the various combinations of NS3 and NS5 or NS5 alone. H is heat control. The phosphorimages were quantified by ImageQuant software and %ssRNA formed was plotted as a function of time (panels B and D).

## Discussion

*Orthoflavivirus* infections pose a significant threat to the U.S. and global public health (46). Prominent members of this genus, including WNV, ZIKV, and DENV, are responsible for recurrent outbreaks of mosquito-borne disease (6, 46). Although most infections are mild and self-limiting, a subset can progress to severe outcomes such as fever, joint and muscle pain, meningitis, encephalitis, paralysis, or death (46). Consequently, there is a pressing need to develop antiviral therapeutics to treat orthoflavivirus infections (47, 48). Orthoflaviviruses are positive-sense, single-stranded RNA viruses that encode two essential enzymes directly responsible for viral RNA replication (6, 7, 11), the NS3 protease–helicase and the NS5 methyltransferase–polymerase, both of which represent tractable targets for antiviral drug discovery (48). The goal of this study was to establish a biochemical framework for understanding orthoflavivirus NS3 helicase activity, how this differs from HCV NS3, and to evaluate if NS5 influences NS3 dsRNA unwinding activity. The work presented here defines fundamental mechanistic distinctions between the orthoflavivirus NS3 helicases and the prototypical HCV NS3 enzyme and establishes a biochemical rationale for functional coupling between NS3 and NS5 during dsRNA unwinding.

Orthoflavivirus NS3 exhibits the canonical SF2 helicase fold and retains RNA-stimulated ATPase activity (**Fig.1**) (13, 15, 20, 49). Strand separation requires ATP hydrolysis (**Fig. 7**), yet NS3 does not form a pre-assembled complex with dsRNA substrate in the absence of ATP-Mg^2+^ (**Fig. 6**). The inability of WNV NS3 to unwind dsRNA under conditions that support robust HCV NS3 activity underscores fundamental differences in the reaction mechanism (**Fig. 2**). HCV NS3 can preassemble on an RNA substrate in the absence of ATP–Mg²⁺, forming a stable complex that produces a rapid burst of unwinding upon addition of ATP–Mg²⁺ (**Fig. 2**) (19). In contrast, orthoflavivirus NS3 required the coincident presence of low concentrations of ATP–Mg²⁺ at the moment of RNA engagement to achieve maximum unwinding (**Fig. 3**, **Fig.6, and Fig. 7**). This observation implies a tightly coupled allosteric linkage between the ATP- and RNA-binding sites of flavivirus NS3. In the absence of nucleotide, NS3 likely resides in an open conformation that fails to engage duplex RNA productively. Binding of ATP–Mg²⁺ may induce domain closure, establishing the interdomain contacts necessary for formation of an active translocation intermediate (12, 14, 50–52). This requirement for nucleotide-dependent assembly rationalizes the sensitivity of unwinding to ATP and Mg²⁺ concentration and suggests that ATP binding and hydrolysis drive not only strand separation but also the formation of the catalytically competent RNA-bound state. Similar behavior has been observed for other helicases whose substrate recognition is dynamically modulated by nucleotide state, implying that orthoflavivirus NS3 likely alternates between low- and high-affinity RNA-binding states during the hydrolysis cycle (53–58). The requirement for an optimal ratio of ATP:Mg²⁺ of 2:0.5 for orthoflavivirus NS3 could suggest that there is tight regulation to coupling ATP hydrolysis to RNA unwinding. Mg²⁺ neutralizes the charges on the β- and γ-phosphates and aligns the ATP in the active site for nucleophilic attack and hydrolysis(14, 50). Excess Mg^2+^ may bind to the protein or ATP in an unproductive form. The presence of excess ATP ensures forward motion of continuous catalysis. HCV NS3 may be more tolerant to higher concentrations of ATP:Mg^2+^ but require the ratio to be near 1. Orthoflavivirus NS3 also exhibits RNA triphosphatase (RTPase) activity which is required to process the 5’-triphosphorylated end prior to capping (6, 7, 10, 22, 23, 39). The HCV NS3 does not process the 5’-end using RTPase activity as the 5’-end of HCV genome is not capped. Notably, the alphavirus genome is also capped and the encoded nsP2 helicase also catalyzes the first step in capping utilizing RTPase activity (59, 60). The nsP2 helicase also has a strict requirement of ATP:Mg²⁺ of 1:0.5 (61). It is interesting to speculate that this optimal ratio exists to tune and regulate the RTPase and ATPase activities independently for specific functions, processing the 5’-end or unwinding dsRNA during viral replication.

The sensitivity of RNA binding to ATP–Mg²⁺ concentration (**Fig. 4**), combined with the requirement for enzyme concentrations exceeding the apparent *K*d,app for RNA (**Fig. 5**), indicates that cooperative interactions or oligomerization may be required for efficient unwinding. It is not uncommon for SF1 or SF2 helicases to have functional cooperativity for unwinding dsRNA (15, 49, 55). This interpretation aligns with prior structural evidence that the NS3 helicase can form higher-order assemblies on RNA and may undergo conformational coupling between adjacent monomers (28, 31, 62, 63). The preference for substrates containing a 5′-single-stranded overhang further suggests that NS3 loading and translocation require specific RNA geometries that orient the helicase along the RNA in a defined 3′→5′ direction. Our results further demonstrate that the orthoflavivirus NS3 helicase exhibits a strong preference for duplex substrates containing a 5′-single-stranded tail on the loading strand (**Fig. 8**). This architecture likely positions NS3 to initiate translocation in the 3′→5′ direction and to maintain continuous contact with the RNA throughout strand displacement (**Fig. 8**). In this context, the substrate dependence of NS3 mirrors the polarity constraints of other SF2 helicases (15, 49), reinforcing the notion that NS3 must engage an extended RNA tract to establish a productive unwinding trajectory.

A key insight from this study is that orthoflavivirus NS5 markedly enhances NS3-mediated unwinding, and this effect is specific to NS3–NS5 pairs (**Fig. 10-12**). This specificity implies a co-evolved interface that enables NS5 to stabilize NS3 in an RNA-engaged conformation or to coordinate their activities during duplex separation and RNA synthesis. Structural and biochemical data support functional interactions between NS3 and NS5 to modulate their activities (10, 11, 38–45). The inability of non-orthoflaviviral polymerases to stimulate NS3 underscores that this interaction reflects a precise mechanistic partnership rather than a generic allosteric effect of RNA-binding proteins (**Fig. 11-12**). We propose that NS5 binding stabilizes an RNA-engaged conformation of NS3, possibly by restricting interdomain flexibility or preventing premature dissociation from the RNA duplex. Such stabilization could increase the dwell time of NS3 on RNA, effectively converting a distributive helicase into a processive motor within the context of the replication complex (10, 11).

The functional coupling of NS3 and NS5 observed here supports a model in which RNA synthesis and unwinding are synchronized events. Within the replication organelle, NS5-catalyzed RNA synthesis would generate RNA duplexes that must be rapidly resolved by NS3. Direct interaction between the two enzymes may coordinate these processes temporally, ensuring that strand displacement and nucleotide incorporation proceed in phase. This level of coordination would parallel the mechanochemical coupling observed in polymerase–helicase assemblies from other RNA viruses and may represent a conserved strategy for regulating replicase efficiency and fidelity.

Together, our results highlight mechanistic principles that distinguish orthoflavivirus NS3 from its HCV counterpart and point toward an essential role for NS5 in helicase function. Future structural and kinetic studies will be required to define how NS5 binding alters NS3 conformational dynamics and whether nucleotide exchange within NS3 is synchronized with polymerase catalysis. Reconstitution of higher-order replication complexes, perhaps within synthetic scaffolds, will enable direct observation of how these enzymes cooperate to remodel RNA during replication. Defining this interplay at molecular resolution will be crucial for understanding how orthoflavivirus replication is regulated and for identifying new strategies to disrupt the viral replicase.

### Experimental procedures

#### Materials

RNA oligonucleotides were from Horizon Discovery Ltd. (Dharmacon). T4 polynucleotide kinase was from ThermoFisher. [γ-^32^P]ATP (6,000 Ci/mmol), [α-^32^P]ATP (3,000 Ci/mmol) were from Revvity. ATP (ultrapure solution) was from Cytiva. NS5’s and PV 3D proteins were produced as described previously (64). All other reagents were of the highest grade available from MilliporeSigma, VWR, or Fisher Scientific.

#### Construction of orthoflavivirus NS3 bacterial expression plasmids

The orthoflavivirus NS3 gene was cloned into the pSUMO bacterial expression plasmid. This system allows for the production of SUMO fusion proteins containing an amino-terminal hexahistidine tag fused to SUMO that can be purified by nickel nitriloacetic acid (Ni-NTA) chromatography and subsequently processed by the SUMO protease, ubiquitin-like-specific protease-1 (Ulp1). The NS3 sequences were derived from the following: WNV NY99 strain (AF404756) and ZIKV strain BeH815744 (AMA12087.1). The NS3 coding region was amplified by PCR and the PCR product was gel purified and cloned into the pSUMO plasmid using BsaI and SalI sites. The final construct was confirmed by sequencing at the Pennsylvania State University’s Nucleic Acid Facility. The D285A mutation was introduced by Quickchange-site-directed mutagenesis.

#### Expression of orthoflavivirus NS3

E. coli Rosetta (DE3) cells were transformed with the pSUMO-NS3 plasmid for protein expression. Rosetta (DE3) cells containing the pSUMO-NS3 plasmid were grown in 100 mL of media (NZCYM) supplemented with 25 μg/mL kanamycin (K25) and 20 μg/mL chloramphenicol (C20) at 37°C until an OD600 of 1.0 was reached. This culture was then used to inoculate 1 L of K75, C60-supplemented ZYP-5052 auto-induction media to an OD600= 0.1. The cells were grown at 37°C to an OD600 of 0.8 to 1.0, cooled to 20°C and then grown for 24 h. Typically, after 24 h at 20°C the OD600 reached ∼10-15. Cells were harvested by centrifugation (6000 x g, 10 min) and the cell pellet was washed once in 200 mL of TE buffer (10 mM Tris, 1 mM EDTA), centrifuged again, and the cell paste weighed. The cells were then frozen and stored at −80°C until used. Frozen cells were thawed and suspended in lysis buffer (20 mM Potassium Phosphate pH 8.0, 20% glycerol, 5 mM Imidazole, 500 mM NaCl, 5 mM BME, 1.4 µg/ml Pepstatin A, and 1.0 µg/ml Leupeptin), 5 mL/g cell pellet. The cell suspension was lysed by passing through a French press (SLM-AMINCO) at 15,000 psi. After lysis, phenylmethylsulfonylfluoride (PMSF) and NP-40 were added to a final concentration of 1 mM and 0.1% (v/v), respectively. While stirring the lysate, polyethylenimine (PEI) was slowly added to a final concentration of 0.25% (v/v) and then centrifuged at 75,000 x g for 30 min at 4 °C. The PEI supernatant was decanted to a fresh beaker, and while stirring, pulverized ammonium sulfate was slowly added to 40% (w/v) saturation. This supernatant was stirred for 30 min after the last addition of ammonium sulfate, and centrifuged at 75,000 x g for 30 min at 4°C. The supernatant was decanted, and the pellet was suspended in buffer B (25 mM HEPES, pH 7.5, 500 mM NaCl, 5 mM BME, 20% glycerol) with 5 mM imidazole. The resuspended ammonium sulfate pellet was loaded onto a Ni-NTA column (Qiagen) at a flow rate of 1 mL/min (approximately 1 mL bed volume/100 mg total protein) equilibrated with buffer B. After loading, the column was washed with twenty column volumes of buffer B containing 5 mM imidazole and five column volumes of buffer B containing 50 mM imidazole. Protein was eluted from the Ni-NTA column with buffer B containing 500 mM imidazole. Fractions were collected and assayed for purity by SDS-PAGE. Ulp1 (1 μg per 10 mg SUMO fusion protein) was added to the purified protein at 4 °C to cleave the SUMO-NS3 fusion protein. The protein was dialyzed overnight in dialysis buffer (20 mM HEPES pH 7.0, 20% glycerol, 10 mM BME, 500 mM NaCl) using a MWCO of 12k-14k dialysis membrane. The protein was removed from dialysis and evaluated by SDS-PAGE to confirm that SUMO fusion was cleaved. The protein was diluted in buffer to 250 mM NaCl final and passed through a Phosphocellulose column. The passthrough was collected, concentrated using Vivaspin turbo 15 columns (10,000 kDa MWCO) and then loaded onto a Cytiva HiLoad Superdex 16/600 200 pg size exclusion column equilibrated with 20 mM HEPES pH 7.0, 20% glycerol, 10 mM BME, and 500 mM NaCl. NS3 fractions were collected, evaluated by SDS-PAGE for purity, pooled, and concentrated. The protein concentration was determined by measuring the absorbance at 280 nm by using a Nanodrop spectrophotometer and using a calculated molar extinction coefficient and stored at −80 °C.

#### NS3 ATPase activity

Reactions were performed at 37 °C, 30 °C and 25 °C. Reactions contained 20 mM MOPS pH 7.0, 50 mM NaCl, 1 mM TCEP, 0.5 mM MgCl_2_, 2 mM ATP and 0.1 μCi/μL [α32P]-ATP. ATP and MgCl_2_ were adjusted accordingly as detailed in results, figures, and legends. Reactions were quenched at various times by addition of EDTA to 10 mM. NS3 was diluted immediately prior to use in 25 mM HEPES pH 7.5, 1 mM TCEP and 20% glycerol. The volume of enzyme added to any reaction was always less than or equal to one–tenth the total volume.

#### Product analysis: TLC

1 µl of the quenched reaction was spotted onto polyethyleneimine-cellulose TLC plates (EM Science). TLC plates were developed in 0.3 M potassium phosphate, pH 7.0, dried and exposed to a PhosphorImager screen. TLC plates were visualized by using a PhosphorImager and quantified by using the ImageQuant software (molecular dynamics) to determine the amount of ATP hydrolyzed to ADP. The amount of ADP was plotted as a function of time.

#### 5’-^32^P-labeling of RNA substrates

RNA oligonucleotides were end-labeled by using [*γ*-^32^P]ATP and T4 polynucleotide kinase. Reaction mixtures, with a typical volume of 50 μL, contained 0.5 μM [*γ*-^32^P]ATP, 1 μM RNA oligonucleotide, 1X kinase buffer, and 0.4 unit/*μ*L T4 polynucleotide kinase. Reaction mixtures were incubated at 37 °C for 60 min and then held at 65 °C for 5 min to heat inactivate T4 PNK.

#### Annealing of dsRNA substrates

dsRNA substrates were produced by annealing 1 μM RNA oligonucleotides in 25 mM HEPES pH 7.5 and 50 mM NaCl in a Progene Thermocycler (Techne). Annealing reaction mixtures were heated to 90 °C for 1 min and slowly cooled (5 °C/min) to 10 °C.

#### RNA binding fluorescence polarization experiments

Increasing concentrations of NS3 were mixed with 1 nM of fluorescein-labeled RNA in a binding buffer containing 20 mM MOPS pH 7.0, 50 mM NaCl, 1 mM TCEP, 0.5 mM MgCl_2_, and 2 mM ATP. ATP and MgCl_2_ were adjusted accordingly as detailed in results, figures, tables, legends. Reactions were incubated at room temperature for 10 minutes in a 384-well non-binding plate and then read with a dual wavelength Ex: 485/20 Em: 528/20 fluorescence polarization cube using a Biotek Synergy H1 plate reader. Data from protein titration experiments were fit to a hyperbola (Eq. 1):

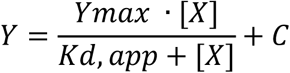

where X is the concentration of protein, Y is degree of polarization, *K*d,app is the apparent dissociation constant, and Ymax is the maximum value of Y.

#### NS3 helicase assay

NS3 was mixed with 5 or 10 nM [^32^P]-labeled unwinding substrate in 20 mM MOPS pH 7.0, 50 mM NaCl, 1 mM TCEP, 0.5 mM MgCl_2_, 2 mM ATP and 0.5 µM DNA Trap. ATP and MgCl_2_ were adjusted accordingly as detailed in results, figures, and legends. The trapping strand is a 9-mer DNA that is complementary to the ^32^P-labeled displaced strand. Reactions were quenched at various times by addition 2X Quench Buffer (200 mM EDTA, 0.66% SDS, 2 mM (UMP) poly(rU), 0.05% Bromophenol Blue, 0.05% Xylene Cyanol, and 10% Glycerol). Specific concentrations of substrate or enzyme, along with any deviations from the above, are indicated in the appropriate legend. NS3 was diluted immediately prior to use in 25 mM HEPES pH 7.5, 1 mM TCEP and 20% glycerol. The volume of enzyme added to any reaction was always less than or equal to one–tenth the total volume. Products were resolved by native PAGE.

#### Product analysis: native PAGE

10 µl of the quenched reaction mixtures was loaded on a 20% acrylamide, 0.53% bisacrylamide native polyacrylamide gel containing 1X TBE. Electrophoresis was performed in 1X TBE at 15 mA. Gels were visualized by using a PhosphorImager and quantitated by using the ImageQuant software (Molecular Dynamics) to determine the amount of strand separation. The fraction of RNA that had been unwound by the helicase was corrected for the efficiency of the trapping strand in preventing reannealing of products. This correction factor was determined for each experiment by heating a sample of the RNA substrate at 95°C for 1 min then cooled to 10°C at a rate of 5°C/min in the presence of the trapping strand, this is indicated as heat control in the appropriate figure (H). The quantity of unwinding substrate that was prevented from reannealing was typically ∼98%. The amount of RNA unwound was plotted as a function of time.

#### Data analysis

All gels shown are representative, single experiments that have been performed at least three to four individual times to define the concentration or time range shown with similar results. In all cases, values for parameters measured during individual trials were within the limits of the error reported for the final experiments. Data were fit by either linear or nonlinear regression using the program GraphPad Prism (GraphPad Software Inc.) or Kaleidagraph (Synergy Software).

## Data availability

All data are incorporated into the article. Constructs and data sets presented in this study are available upon request.

## Acknowledgements

We thank Jacob Perryman and Rajni Sharma for the contributions to the expression and purification flavivirus NS3 enzymes.

## Author contributions

**Jamie J. Arnold:** Conceptualization, Methodology, Formal analysis, Investigation, Resources, Data Curation, Writing - Original Draft, Writing - Review & Editing, Visualization, Funding Acquisition; **Shubeena Chib:** Conceptualization, Methodology, Formal analysis, Investigation, Resources, Data Curation, Writing - Original Draft, Writing - Review & Editing, Visualization; **Craig E. Cameron:** Conceptualization, Formal analysis, Writing - Review & Editing, Supervision, Project administration, Funding Acquisition.

## Funding and additional information

This work was supported by the National Institutes of Health [R01AI045818 to C.E.C., J.J.A.]

## Abbreviations

ADP: adenosine diphosphate
ATP: adenosine triphosphate
BME: β-mercaptoethanol
DENV: dengue virus
dsRNA: double-stranded RNA
EDTA: ethylenediaminetetraacetic acid
HCV: hepatitis C virus
HEPES: 4-(2-hydroxyethyl)-1-piperazineethanesulfonic acid
MOPS: 3-(N-morpholino)propanesulfonic acid
MWCO: molecular weight cut-off
Ni-NTA: nickel–nitrilotriacetic acid
NS3: non-structural protein 3
NS5: non-structural protein 5
nt: nucleotide(s)
NTPase: nucleoside triphosphatase
PAGE: polyacrylamide gel electrophoresis
PCR: polymerase chain reaction
PEI: polyethylenimine
PNK: polynucleotide kinase
PV: poliovirus
RdRp: RNA-dependent RNA polymerase
RTPase: RNA triphosphatase
SDS-PAGE: sodium dodecyl sulfate–polyacrylamide gel electrophoresis
SF2: superfamily 2 helicase
ssRNA: single-stranded RNA
SUMO: small ubiquitin-like modifier
TCEP: tris(2-carboxyethyl)phosphine
TLC: thin-layer chromatography
Ulp1: ubiquitin-like-specific protease 1
WNV: West Nile virus
WT: wild type
ZIKV: Zika virus

